# Activity/rest rhythms of *Drosophila* populations selected for divergent eclosion timing under temperature cues

**DOI:** 10.1101/831347

**Authors:** Lakshman Abhilash, Kalliyil Arshad, Vasu Sheeba

**Author notes:** Correspondence: Vasu Sheeba, Behavioural Neurogenetics Laboratory, Neuroscience Unit, Jawaharlal Nehru Centre for Advanced Scientific Research, Jakkur, Bangalore – 560064, Karnataka, India.

## Abstract

Despite the fact that the rhythm in adult emergence and rhythm in locomotor activity are two different rhythmic phenomena that occur at distinct life-stages of the fly life cycle, previous studies have hinted at similarities in certain aspects of the organisation of the circadian clock driving these two rhythms. In an earlier study, we have shown that selection on timing of adult emergence behaviour in populations of *Drosophila melanogaster* leads to the co-evolution of temperature sensitivity of circadian clocks driving eclosion. In this study, we were interested in asking if temperature sensitivity of the locomotor activity rhythm has evolved in our populations with divergent timing of adult emergence rhythm, with the goal of understanding the extent of similarity (or lack of it) in circadian organisation between the two rhythms. We found that in response to simulated jetlag with temperature cycles, *late* chronotypes (populations selected for predominant emergence during dusk) indeed re-entrain faster than *early* chronotypes (populations selected for predominant emergence during dawn), thereby indicating enhanced sensitivity of the activity/rest clock to temperature cues in these stocks (entrainment is the synchronisation of internal rhythms to cyclic environmental time-cues). Additionally, we found that *late* chronotypes show higher plasticity of phases across regimes, day-to-day stability in phases and amplitude of entrainment, all indicative of enhanced temperature sensitive activity/rest rhythms. Our results highlight remarkably similar organisation principles between emergence and activity/rest rhythms.

## Introduction

Many insects show rhythmic emergence of adults from their pupal cases. These rhythms persist in the wild and in the laboratory under both light/dark (LD) cycles and constant conditions (Saunders, 2002). Among them, a large proportion of species time this remarkable phenomenon to occur just after dawn. These include, for instance, the yellow dung fly (*Scopeuma stercoraria*), the Queensland fruit-fly (*Dacus tryoni*), moths (*Pectinophora gossypiella* and *Heliothis zea*) and many *Drosophila* species (Saunders, 2002). These observations have raised two interesting questions. What is the mechanism by which organisms restrict emergence to certain times of the day? Why is emergence predominantly restricted to dawn in so many insect species? These questions have been of interest for many decades, and we are now aware of the presence of biological time-keeping mechanisms (also referred to as circadian clocks) that generate and drive rhythmicity in many aspects of behaviour and, across almost all living beings (Dunlap et al., 2004). To the question of why emergence is restricted to dawn, Colin S Pittendrigh in the mid-1950s hypothesised that organisms must have evolved to time emergence to the time of the day when humidity is high and temperature is low. This was thought to allow efficient wing expansion in pharate adults and therefore enable survival (Pittendrigh, 1954).

Pittendrigh’s hypothesis implied that timing of emergence and modalities that allow sensation of and responses to temperature and/or humidity are intimately linked. Adaptations to capitalise on a temporal niche, in many other organisms, are also thought to be multi-tiered such that multiple aspects of behaviour, physiology and morphology evolve together (Daan, 1981). For instance, in addition to circadian clock properties, waxy cuticles to prevent water loss and enhanced vision are thought to have evolved in diurnal insects, while improved sound and olfaction are thought to have evolved in nocturnal birds and mammals (Daan, 1981). However, clear demonstration of the genetic association of various aspects ofphysiology and circadian clock properties via the evolution of behavioural timing has been lacking. With the goal of understanding such relationships, our laboratory generated and currently maintains four large and outbreeding *Drosophila melanogaster* populations that are laboratory (artificially) selected for morning and evening adult emergence (first described in Kumar et al., 2007).

In relation to Pittendrigh’s 1954 hypothesis, we have recently demonstrated that laboratory selection for evening timing of emergence (as opposed to morning) is strongly associated with the co-evolution of enhanced temperature sensitivity of the circadian clock circuit regulating adult emergence rhythms (Abhilash et al., 2019). This clearly demonstrates a genetic correlation between behavioural phase and temperature responses, and is in agreement with the above hypothesis. Although Pittendrigh’s argument was made based on cycles of both temperature and humidity, due to technical limitations, there are barely any studies on the role of humidity in emergence rhythms. Effects of temperature, on the other hand, are fairly well studied in both adult emergence and locomotor activity rhythms (Das and Sheeba, 2017; Pittendrigh, 1954; Selcho et al., 2017).

It is important to note here that i) while the emergence rhythm is a population level phenomenon (each individual can emerge from its pupal case only once), locomotor activity is an individual level rhythm, ii) further, adult emergence and adult locomotor activity are two very different physiological processes occurring at two entirely different life-stages of the fly life cycle. Despite that, interestingly, the first study that isolated and described the effects of *period* (*per*) mutation on behavioural rhythms in *Drosophila*, found that both emergence and activity/rest rhythms are affected in a similar manner for all alleles of the *per* locus (Konopka and Benzer, 1971). Subsequently, it was demonstrated that the small ventral lateral neurons (s-LNvs) are necessary to drive behavioural rhythms in eclosion and locomotor activity under constant conditions (Grima et al., 2004; Myers et al., 2003; Stoleru et al., 2004), thereby illustrating the close overlap in the timing machinery regulating both rhythms. Additional evidence also comes from the fact that the free-running period (FRP) estimated using eclosion and activity/rest rhythms are strongly positively correlated with each other in flies from our morning and evening selected populations (Kumar et al., 2007; Nikhil, Abhilash, et al., 2016), again highlighting a common machinery regulating both behaviours. In light of these similarities in organisation of the circadian clock across rhythms spanning two very different behaviours, we asked if temperature sensitivity of the clock regulating the adult locomotor activity rhythm also evolved in the evening emerging flies? In reference to the way the circadian network is organised, what does this imply?

To address this question, we first subjected our flies to simulated jetlag of 6-h phase advance (equivalent to eastward travel, egs., London to Bangkok) and 6-h phase delay (equivalent to westward travel, egs., London to Chicago) using temperature cycles alone. We found that flies from the evening populations resynchronise to phase-shifted temperature cycles much faster than the morning and *control* populations. This result indicated differences in the temperature sensitive components of the circadian circuit in the morning and evening flies. To further understand the nature of differences in sensitivity, we explored their behaviour under temperature cycles with different durations of warm phase under otherwise constant darkness. We analysed various aspects of the activity/rest rhythm – phase, accuracy (day-to-day variation in phases), power of the rhythm (see material and methods) and consolidation of rhythm under entrainment. We also examined period of the rhythm and its amplitude under constant darkness post entrainment to the aforementioned temperature cycles. Subsequently, we also analysed the behaviour of these flies under light/dark cycles at two different constant ambient temperatures to understand degree of waveform plasticity under different constant temperatures. Our results suggest that selection for divergent timing of emergence behaviour is also associated with increased temperature sensitivity of the clock regulating activity/rest rhythms. Interestingly, we find that this increased sensitivity is brought about predominantly by the evening bout of activity.

## Materials and Methods

### Selection protocol and fly husbandry

All experiments reported in this manuscript made use of 4 sets of populations each – *early*_*(1-4)*_, *control*_*(1-4)*_ and *late*_*(1-4)*_ that have been derived in our laboratory through artificial selection for morning eclosion (*early*) and evening eclosion (*late*). Each of the four sets have independent genetic architecture and are referred to as ‘blocks’. Selection was induced in each of these sets separately and simultaneously. Every generation ~300 eggs are collected and dispensed into vials that are then maintained in a light, temperature and humidity controlled cubicle under LD 12:12 (light/dark cycles with 12-h light and 12-h darkness) at 25±0.5 °C and 65±5% RH. On the 9^th^ – 13^th^ day after egg collection, only flies emerging in a four hour window starting 3 hours before lights-ON and ending an hour after lights-ON (or ZT21-ZT01, where ZT00 is zeitgeber time 00 and refers to the time at which lights come ON) are collected to form the breeding pool for the next generation of the *early* flies. Similarly, all the flies emerging on the same days but between ZT09 and ZT13 are collected to form the breeding pool of the next generation of *late* flies. For *control* populations, flies emerging throughout the day are collected to form the next generation. All adult flies are maintained in Plexiglas cages with culture medium in petriplates at a roughly 1:1 sex ratio and adult density of ~1500 flies per cage. On the 18^th^ day after the previous egg collection, flies in the cages are provided food supplemented with live yeast paste (as a protein supplement) and on the 21^st^ day, eggs are collected in the same manner described above for the new generation. Therefore, our flies are maintained on a 21-day discrete generation cycle and all flies used in the experiments reported here have undergone over 280 (~16 years) generations of selection. All experiments were performed on progeny of flies that experienced one generation of common rearing, to avoid maternal effects on traits being measured which may confound our interpretations (Bonduriansky and Day, 2009).

### Behavioural experiments

~300 eggs from each of the 12 populations were collected (as during maintenance) and dispensed into 5-10 vials and maintained under standard maintenance regime. Two sets of thirty two 3-5 day old virgin males were collected, and under minimal CO_2_ anaesthesia were transferred to 5mm locomotor tubes. These sets were then recorded using the *Drosophila* Activity Monitor (DAM) system under warm:cold cycles, TC 12:12 (thermophase: 28 °C; cryophase: 19 °C) for 4-5 cycles before simulating the jetlag. One of these sets was given a 6-h advance phase shift, while the other set was given a 6-h delay phase shift for 10 cycles before all the flies were transferred to constant darkness at 19 °C for a few cycles to judge phase control (an essential property of circadian clocks wherein phase of activity on first day in constant darkness and temperature continues from the phase on last day of temperature cycles).

In the second batch of experiments, three sets of thirty two 3-5 day old virgin male flies were collected in the same manner as described in the previous paragraph. One set each was then subjected to TC 06:18, TC 12:12 and TC 18:06, under constant darkness for 7 days (thermophase: 28 °C; cryophase: 19 °C). On the 8^th^ day, flies transitioned from TC to constant cryophase of 19 °C for 6-8 days, so as to enable analyses of FRP.

In the third batch of experiments, two sets of thirty two 3-5 day old virgin males were collected as described above. One set was recorded under LD 12:12 (~70 lux during the photophase) at 19 °C and the other at 28 °C. Both these sets were recorded under their respective conditions for 7-8 days, before being transferred to DD (constant darkness) under their respective constant temperatures. Flies were maintained under free-running conditions for 6 days, so as to allow estimation of the FRP. We used FRP data of all our stocks to facilitate comparisons with the experiments reported here from a previous experiment performed by Lakshman Abhilash, entrained behaviour of which is published elsewhere (Nikhil, Abhilash, et al., 2016).

### Data analysis

#### Rates of re-entrainment

To estimate rates of re-entrainment to 6-h advance and delay, we marked phases of offset for each fly on each day for all 12 populations using RhythmicAlly (Abhilash and Sheeba, 2019). We then calculated the daily phase relationship as phase of offset of activity – phase of offset of thermophase. For each pre-jetlag cycle, we calculated the average phase relationship across flies. Then we computed the average inter-individual variation in between-fly phase relationships. Subsequently, we multiplied this measure by 1.96 to get a 95% confidence band around the mean inter-cycle phase relationship. We did this for all populations and then examined the dynamics of phase relationship change across days for each fly. A fly was considered re-entrained when its phase relationship re-entered the confidence band and stayed inside the band for at least two subsequent cycles. The number of cycles taken for each fly to re-entrain was used as a measure of number of transients taken for re-entrainment. These values were averaged across flies for obtaining block means. Two separate two factor mixed model randomised block design ANOVAs were used to analyse the effect of selection on number of transients taken for re-entrainment, each for the 6-h advance regime and 6-h delay regime. Selection was used as a fixed factor and block as a random factor.

#### Activity profiles under TC cycles

We analysed the activity profiles for all populations under TC 06:18, TC 12:12 and TC 18:06. Raw DAM data was scanned and monitor files were saved in 20-min bins. These data files were analysed using RhythmicAlly (Abhilash and Sheeba, 2019). Individual profiles were downloaded and were re-organised to 1-h bins. Activity counts were then averaged across flies within each block to obtain population-wise profiles. As on multiple previous occasions (e.g., Nikhil et al., 2014; Srivastava et al., 2019), Centre of Mass (CoM) was used as a non-subjective phase marker (*ψ*_*CoM*_) so that changes in the waveform under different regimes are captured reliably. Owing to the bimodality of activity profiles under TC 12:12 and TC 18:06, an angle doubling transformation was performed before computing *ψ*_*CoM*_ (Batschelet, 1981). For TC 06:18, *ψ*_*CoM*_ was computed without the angle doubling transformation. FRP of flies experiencing constant conditions post aforementioned entrainment regimes was quantified using the chi-square periodogram implemented in RhythmicAlly (Abhilash and Sheeba, 2019).

We used circular *r* as a proxy measure of normalised amplitude (owing to its significance in describing the consolidation of a peak), as has been used before (Abhilash et al., 2019). Similar to the computation of phase, angle doubling was performed on activity profiles under TC 12:12 and 18:06 only. Intrinsic amplitude of each of these stocks were estimated using ActogramJ (Schmid et al., 2011). First, average actograms for each block was generated using data post entrainment to their respective temperature cycles. Then a chi-squared periodogram analysis was done on each population to estimate the average FRP. Then activity profile was generated using modulo-FRP for each block. Amplitude was measured as the maximum activity count – minimum activity count in each of these profiles. To estimate accuracy (cycle to cycle variation in entrained phase), we calculated *ψ*_*CoM*_ for each fly and each cycle during entrainment. Accuracy was defined as inverse of standard deviation in *ψ*_*CoM*_ across cycles for each fly. These values were then averaged across flies to obtain block means.

Phases, FRP, intrinsic amplitude, amplitude and accuracy of entrainment were analysed using two factor mixed model randomised block design ANOVAs, wherein selection was treated as a fixed factor and block as a random factor.

#### Activity profiles under constant ambient temperature regimes

To examine behaviour under LD 12:12 at different constant ambient temperatures, average profiles were obtained as described above. Using the population-wise profile data we calculated a ratio of total activity during the day to the total activity during the night for each population (day/night ratio). These were quantified for profiles under both temperatures. Further, we were interested in asking if anything about the waveform in different temperatures changed differently across populations. For this, we used the 1-h binned activity profiles and computed difference in activity level at each time-point between two temperatures. This difference was squared and the sum of these squared differences (SSD) across the entire cycle was calculated as a measure of deviance of rhythm waveform in the two temperatures. From the 1-h binned profiles, we also computed total activity in a morning window (ZT01-06) and an evening window (ZT06-11) to assess the individual contributions of morning and evening bouts of activity to potential differences in temperature sensitivity. FRP for all these flies under constant darkness at 19 and 28 °C were assessed in RhythmicAlly (Abhilash and Sheeba, 2019) using the chi-squared periodogram. Similar analyses were performed to estimate FRP of flies under DD at 25 °C.

The day/night ratio, total morning, total evening activity counts and FRP post entrainment to LD cycles under different temperatures were analysed using three-factor mixed model randomised block design ANOVAs using block means. In these ANOVAs selection and temperature were treated as fixed factors and block was treated as a random factor. The SSD values were analysed using a two-factor mixed model randomised block design ANOVA wherein selection was used as a fixed factor and block was used as a random factor. All statistical analyses were followed by a Tukey’s Honestly Significant Difference (HSD) post-hoc test, to generate error bars that facilitate easy visual hypothesis testing. All results were considered significant at *α* = 0.05.

## >Results

### Re-entrainment to simulated jetlag

As a first step to understand if our *early* and *late* flies differ in sensitivity of their circadian clocks to temperature, we analysed their behaviours to simulated jetlag of 6-h phase advance and 6-h phase delay under TC 12:12. From the representative actograms, one can see that when flies are subjected to a 6-h phase advance regime, all three stocks resynchronise to the phase-shifted TC cycles fairly quickly and take about the same time (Fig. 1A-top). On the other hand it appears that all stocks take much longer to re-entrain to a 6-h phase delay; however the *late* stocks re-entrain sooner than the *early* and *control* stocks (Fig. 1A-bottom). These patterns were clearly visible when we analysed the dynamics of phase-relationships for each stock across days for both phase-shifted regimes (Fig. 1B).

**Figure 1:**
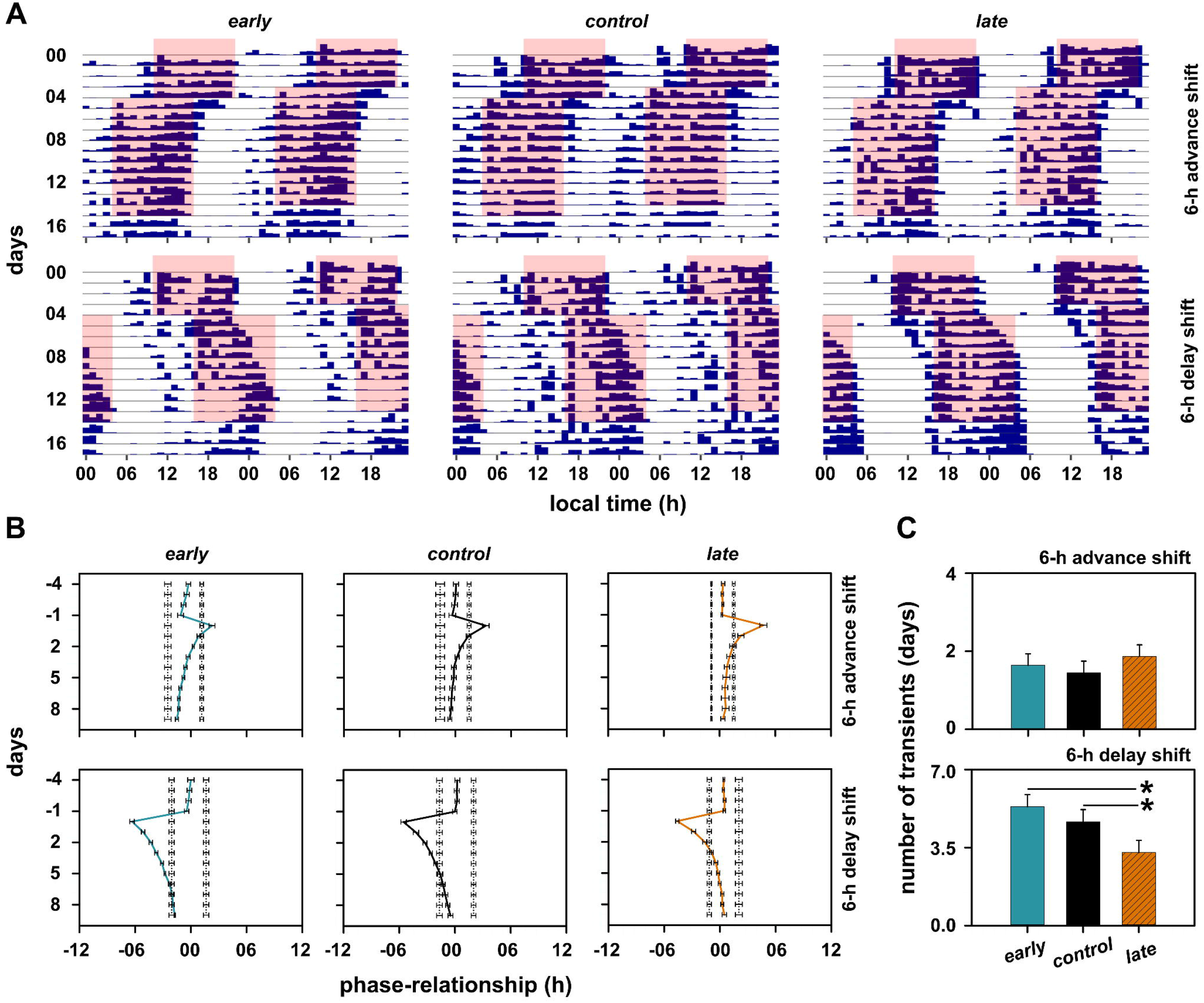
Representative actograms of flies experiencing 6-h advance (A-top) and 6-h delay (A-bottom) shift of TC cycles. Red shaded region indicate the thermophase (28 °C) of the TC cycles (cryophase temperature was 19 °C). Also shown are phase-relationship resetting dynamics averaged over all 4 block for each population under 6-h advance (B-top) and 6-delay (B-bottom) shifts. Error bars here are SEM. Day 0 on the *y*-axis represents the first day of simulated jetlag. Panel (C) shows the average number of transient cycles taken by each population to re-entrain to the 6-h advance (top) and 6-h delay (bottom) shifts. Error bars in this panel are 95% CI from a Tukey’s HSD test at *α* = 0.05. Therefore, means with non-overlapping error bars are statistically significantly different from each other. Additionally, asterisks are drawn to indicate means that are significantly different.

We quantified this and found that all three stocks take fewer than 2 cycles to re-entrain to 6-h phase-advanced TC cycles and there are no between-stock differences in number of transient cycles (*F*_*2,6*_ = 2.37, *p* > 0.05; Figs. 1B-top and 1C-top). On the other hand, under the 6-h phase-delayed TC cycles, while the *early* stocks took about 5.3 cycles and the *control* stocks took ~4.6 cycles, the *late* stocks only took ~3.2 cycles to re-entrain (Figs. 1B-bottom and 1C-bottom). These between stock differences were statistically significant as was revealed by the significant main effect of selection on number of transients (*F*_*2,6*_ = 17.53, *p* < 0.05). Results from this experiment suggest the presence of a circadian clock circuit with ehanced temperature sensitivity in the *late* stocks.

### Activity/rest rhythms under TC cycles (06:18, 12:12, 18:06) and constant conditions post entrainment

To further understand if underlying differences in clock sensitivity to thermal cues contributes to plasticity in phases under different durations of warmth in warm:cold cycles (TC cycles), we examined their behaviour under three different thermoperiods (TC 06:18, TC 12:12 and TC 18:06) under otherwise constant darkness. Across thermoperiods, most activity for both *early* and *late* stocks were restricted to the thermophase, as can be clearly seen in the actograms (Figs. 2A and 2B). Moreover, it appears as though under TC 12:12 and TC 18:06, the *late* chronotypes have delayed evening activity compared to the *early* chronotypes (Figs. 2A-middle and right and 2B-middle and right). Therefore, we quantified entrained phases and found that under TC 06:18 the *ψ*_*CoM*_ was not different between stocks (*F*_*2,6*_ = 4.20, *p* > 0.05; Fig. 2C-top-left), whereas, as expected from the actograms, the *ψ*_*CoM*_ of *late* chronotypes was significantly delayed compared to that of *early* chronotypes under both TC 12:12 (*F*_*2,6*_ = 7.10, *p* < 0.05) and TC 18:06 (*F*_*2,6*_ = 5.91, *p* < 0.05; Fig. 2C-top-middle and top-right).

**Figure 2:**
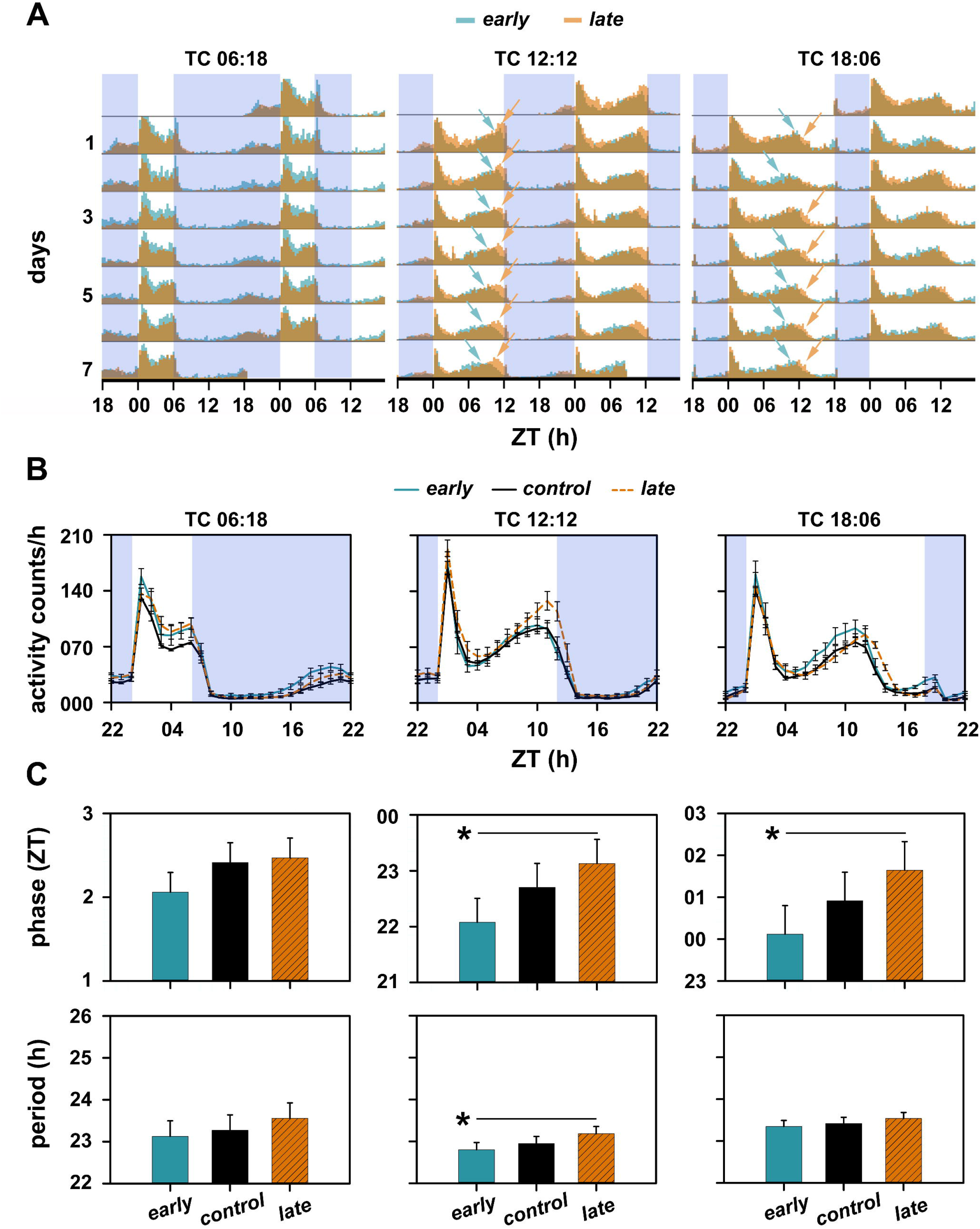
(A) Actograms of *early* and *late* stocks under three different thermoperiods, i.e., TC 06:18 (Thermophase: 6-h, Cryophase: 18-h), TC 12:12 and TC 18:06. Also shown are activity profiles averaged over 4 blocks in panel (B) for all three thermoperiods. Error bars in panel (B) are SEM. The blue shaded region in panels (A) and (B) indicate the cryophase of the TC cycles. (C) Phases of center of mass (*ψ*_*CoM*_) of the three stocks under all three thermoperiods (top panel). Phases for TC 12:12 and TC 18:06 were calculated after an angle doubling transformation was applied, due to the bimodality of the profiles. No such transformation was performed for calculating phase under TC 06:18. Also depicted are free-running periods (FRP) of each stock under constant conditions after being entrained to each of the thermoperiods for ~7 cycles (bottom panel). All error bars in panel (C) are 95% CI from a Tukey’s HSD test at *α* = 0.05. Therefore, means with non-overlapping error bars are statistically significantly different from each other. Additionally, asterisks are drawn to indicate means that are significantly different.

Furthermore, earlier experiments from our laboratory have reported the presence of FRP differences between the stocks under DD at 25 °C (Kumar et al., 2007; Nikhil, Abhilash, et al., 2016). Our results, therefore, implied that entrainment to temperature cycles in these stocks can be explained within the framework of the non-parametric model of entrainment, a central prediction of which is that longer FRP is associated with delayed phase (Pittendrigh and Daan, 1976). To test this, we first analysed FRP values of the flies that experienced the presented TC and were transferred to DD at 19 °C. We found that there was no statistically significant between-stock difference in FRP when flies were transferred to constant conditions after TC 06:18 (*F*_*2,6*_ = 1.70, *p* > 0.05; Fig. 2C-bottom-left; Table 1). Interestingly, after exposure to TC 12:12 the *late* chronotypes showed significantly longer FRP in constant conditions compared to that of the *early* stocks (*F*_*2,6*_ = 5.90, *p* < 0.05; Fig. 2C-bottom-middle; Table 1). However, although there were between stock differences in phase under TC 18:06, there was no between-stock difference in FRP (*F*_*2,6*_ = 2.10, *p* > 0.05; Fig. 2C-bottom-right; Table 1).

**Table 1:** Mean values of FRP post entrainment to LD 12:12 at different constant ambient temperatures and post entrainment to TC cycles with three different thermoperiods ± SEM for all three stocks are reported.

| FRP<br>post entrainment to | *early* | *control* | *late* |
| --- | --- | --- | --- |
| **LD 12:12, 19 °C** | 23.34±0.06-h | 23.62±0.07-h | 23.98±0.08-h |
| **LD 12:12, 25 °C** | 23.64±0.07-h | 23.96±0.07-h | 24.46±0.14-h |
| **LD 12:12, 28 °C** | 23.91±0.05-h | 24.39±0.09-h | 24.88±0.05-h |
| **TC 06:18, DD** | 23.12±0.05-h | 23.27±0.24-h | 23.56±0.09-h |
| **TC 12:12, DD** | 22.80±0.06-h | 22.95±0.10-h | 23.18±0.05-h |
| **TC 18:06, DD** | 23.35±0.07-h | 23.42±0.07-h | 23.54±0.04-h |

Results from both previous sections were suggestive of between-stock differences in phase-dependent sensitivity of the circadian clock to temperature, typically characterised by a phase-response curve (PRC). Therefore, we further analysed other predictions under the assumption of divergent temperature PRCs of the *early* and *late* stocks.

### Robustness, amplitude and accuracy of entrainment under TC cycles

Previous studies have linked divergent PRCs to differences in intrinsic amplitude of the circadian oscillator, its amplitude under entrainment, power of periodogram and accuracy of entrainment (Beersma et al., 1999; Brown et al., 2008; Nikhil, Vaze, et al., 2016; Vitaterna et al., 2006). We examined these properties in flies that were exposed to TC and subsequently placed in DD. Firstly, we found that the *late* stocks had significantly higher amplitude of entrainment, estimated using circular *r* (a measure of how sharp the peak is), under both the asymmetric thermoperiods TC 06:18 (*F*_*2,6*_ = 20.77, *p* < 0.05) and TC 18:06 (*F*_*2,6*_ = 9.17, *p* < 0.05; Fig. 3A-left and right). However, there was no significant between-stock difference in the amplitude under entrainment to TC 12:12 (*F*_*2,6*_ = 4.38, *p* > 0.05; Fig. 3A-middle). Further, there was no significant between-stock difference in intrinsic amplitude under DD post TC 06:18 (*F*_*2,6*_ = 0.67, *p* > 0.05) or post TC 18:06 (*F*_*2,6*_ = 1.26, *p* > 0.05; Fig. 3B-left and right). However, the *late* stocks had significantly higher intrinsic amplitude, post entrainment to TC 12:12 (*F*_*2,6*_ = 9.86, *p* < 0.05; Fig. 3B-middle). These results imply higher amplitude expansion of the *late* stocks under specific TC regimes, thereby suggesting stronger temperature sensitivity in these stocks. We reasoned that periodogram power during entrainment could provide additional evidence for enhanced temperature sensitivity in the *late* stocks. We found that power was significantly higher for *late* stocks only under entrainment to TC 12:12 (*F*_*2,6*_ = 5.83, *p* < 0.05; Fig. 3C). Subsequently, we analysed the accuracy of entrainment under all three regimes. We found that under short thermoperiod there was no between-stock differences in accuracy of entrainment (*F*_*2,6*_ = 1.24, *p* > 0.05; Fig. 3D-left). Under both, TC 12:12 and TC 18:06, there was a significant main effect of selection such that the *late* chronotypes showed significantly increased accuracy of entrainment (TC 12:12: *F*_*2,6*_ = 9.20, *p* < 0.05; TC 18:06: *F*_*2,6*_ = 6.46, *p* < 0.05; Fig. 3D-middle and right).

**Figure 3:**
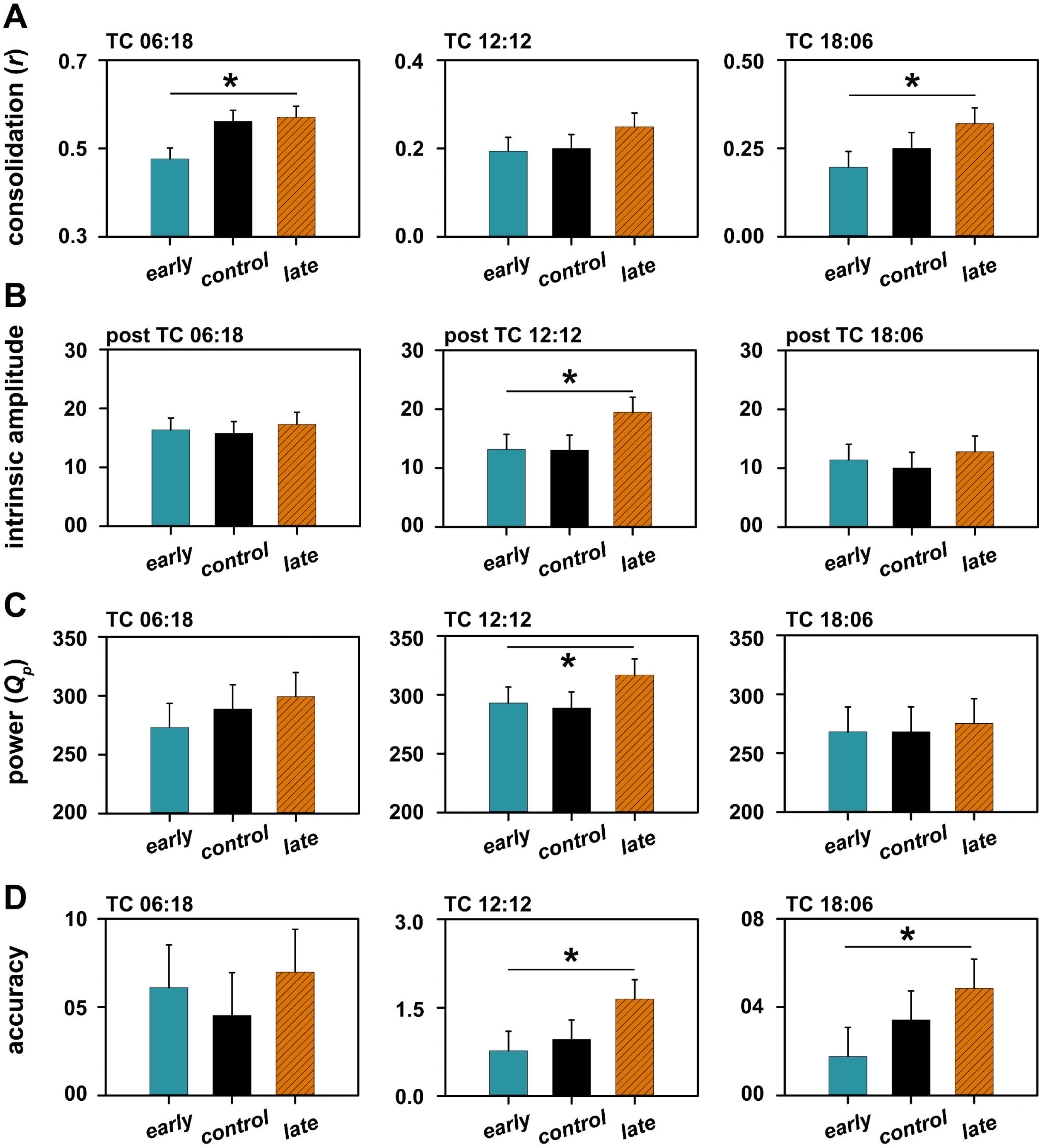
Depicted are consolidation as a measure of amplitude under TC cycles (A), intrinsic amplitude under constant conditions post entrainment (B), power of the chi-square periodogram (C) and accuracy of entrainment (D) for all three stocks under all three thermoperiods. All error bars are 95% CI from a Tukey’s HSD test at *α* = 0.05. Therefore, means with non-overlapping error bars are statistically significantly different from each other. Additionally, asterisks are drawn to indicate means that are significantly different.

In summary, we found evidence for the evolution of temperature sensitivity in the activity/rest rhythms in populations selected for divergent timing of adult emergence rhythm. Additionally, these results also suggest that most features of entrainment can be well explained within the framework of the non-parametric model described above, which makes use of predictions using the FRP and the PRC of the clock.

### Activity/rest rhythms under LD 12:12 and constant ambient temperatures

Owing to the fact that selection for evening emergence contributed to enhanced phase plasticity of emergence rhythms even under LD and different constant ambient temperatures (Abhilash et al., 2019), we next analysed the activity/rest behaviour of our stocks under said regimes. Visual inspection of the activity/rest profiles of *early*, *control* and *late* stocks under 19 °C indicated higher evening activity in the *late* chronotypes (Fig. 4A-left). On the other hand, profiles under 28 °C looked largely similar (Fig. 4A-right), except the slightly increased morning activity in the *late* flies. To quantify this, we calculated the ratio of total day-time activity to total night-time activity (day/night ratio, henceforth). We found that there was a significant main effect of temperature-regime such that there was higher day-time activity under 19 °C as has earlier been shown (*F*_*1,3*_ = 98.10, *p* < 0.05; Majercak et al., 1999). However, using day/night ratio as a measure, we did not find a statistically significant effect of selection × temperature-regime interaction (*F*_*2,6*_ = 0.54, *p* > 0.05; Fig. 4B-left), thereby implying that all three stocks responded similarly to cool and warm ambient temperatures.

**Figure 4:**
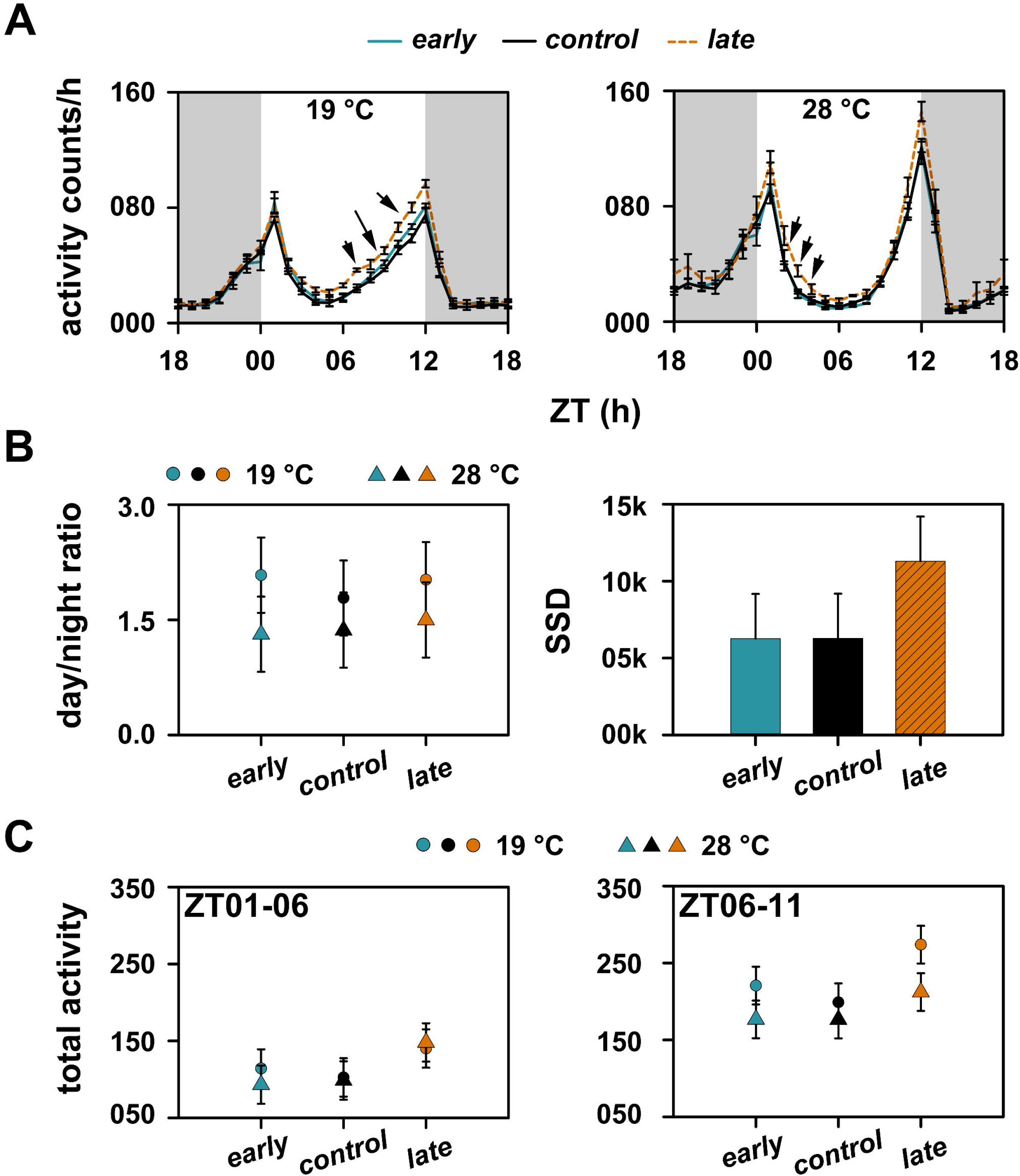
Depicted are locomotor activity profiles of *early*, *control* and *late* stocks under LD 12:12 at 19 °C (A-left) and 28 °C (A-right). The black arrowheads mark parts of the profiles where *late* stocks show higher activity. Gray shaded regions in panel (A) depict the dark phase of the LD cycle. Error bars in this panel are standard error of the mean (SEM). Also shown are day/night ratio of total activity under both temperatures for all three stocks (B-left) and the total sum of square difference between profiles under 19 and 28 °C for all three stocks (B-right). Panel (C) shows total activity in defined first (left) and second (right) halves of the light phase, ignoring the activity during the peak times. All error bars in panels (B) and (C) are 95% CI following a Tukey’s HSD test at *α* = 0.05. Therefore means with non-overlapping error bars are significantly different from each other.

We then examined divergence in waveform across stocks and regimes by calculating SSD (see materials and methods) as a measure of extent of plasticity that *early*, *control* and *late* stocks show in response to different constant ambient temperatures. Although the SSD in *late* stocks is almost twice as much as that in *early* and *control* stocks (Fig. 4B-right), the ANOVA did not detect a main effect of selection, thereby implying that the stocks respond similarly to change in ambient temperature (*F*_*2,3*_ = 4.69, *p* = 0.059).

However, owing to the fact that the SSD difference between stocks was marginally non-significant and that there were trends of between stock differences we quantified total activity in a morning window (ZT01-06) and an evening window (ZT06-11) for all three stocks under 19 and 28 °C. We found that although total morning activity was significantly higher for the *late* stocks (*F*_*2,6*_ = 17.65, *p* < 0.05), there was no selection × temperature-regime interaction (*F*_*2,6*_ = 1.33, *p* > 0.05; Fig. 4C-left), thereby indicating the absence of a stock specific response to temperature. However, in case of total activity in the evening, although there was no statistically significant selection × temperature-regime interaction (*F*_*2,6*_ = 2.56, *p* > 0.05), there is a clear trend of the *late* stocks suppressing activity more strongly under 28 °C relative to the *early* and *control* stocks (Fig. 4C-right). These results are suggestive, although not strongly, of increased plasticity of the activity/rest waveform in the *late* chronotypes in response to different constant ambient temperatures under LD cycles.

### Plasticity of FRP post entrainment to different temperature regimes

While analysing features of the activity/rest rhythm under entrainment, one result that piqued our curiosity was the absolute scale on which FRP varied post entrainment to different thermoperiods (Fig. 2C-bottom; Table 1). While period values ranged between 23.1-h (*early)* to 23.5-h (*late)* stocks and 23.3-h in the *early* stocks to 23.5-h in the *late* stocks post entrainment to TC 06:18 and TC 18:06 (Table 1), respectively, overall period values were much lower, post entrainment to TC 12:12 (Table 1). The period values ranged from 22.8-h in the *early* stocks to 23.2-h in the *late* stocks (see Fig. 2C-bottom; Table 1). Moreover, the absence of statistically significant difference in FRP between *early* and *late* stocks post entrainment to TC 06:18 and 18:06, and the difference under TC 12:12 implies stock specific responses of FRP to cycling temperatures (Table 1).

We, then, examined the FRP of these stocks under DD at 19, 25 and 28 °C post entrainment to LD 12:12 at these respective temperatures. We found that there was a statistically significant main effect of selection such that the *late* stocks had significantly longer FRP than the *early* stocks (*F*_*2,6*_ = 189.70, *p* < 0.05; Fig. 5; Table 1). Also, there was a significant main effect of temperature such that FRP lengthened with increase in temperature (*F*_*2,6*_ = 50.00, *p* < 0.05; Fig. 5; Table 1). Such overcompensation of FRP to changing temperatures in insects is an already established phenomenon (see Saunders, 2002). However, although there was no statistically significant selection × temperature-regime interaction (*F*_*2,6*_ = 1.10, *p* > 0.05), upon closer inspection of the period values some very interesting patterns emerge. On one hand, the FRP of *early* stocks at 19 and 28 °C, although different from each other, do not significantly differ from their value at 25 °C (Fig. 5). On the other hand, the *late* stocks lengthen their FRP at 25 °C compared to its value under 19 °C implying increased responsiveness of the FRP to temperature in these stocks (Fig. 5). Moreover, under 19 °C, the FRP of *early* and *late* stocks, although different from each other, are not different from the FRP of *control* stocks (Fig. 5). The difference in FRP between *early* and *control* stocks only appear under 28 °C, whereas for the *late* stocks the difference is apparent at 25 °C itself, again implying enhanced sensitivity of FRP (with reference to compensatory mechanisms) to temperature in the *late* stocks (Fig. 5). However, there is only weak statistical evidence in favour of these ideas.

**Figure 5:**
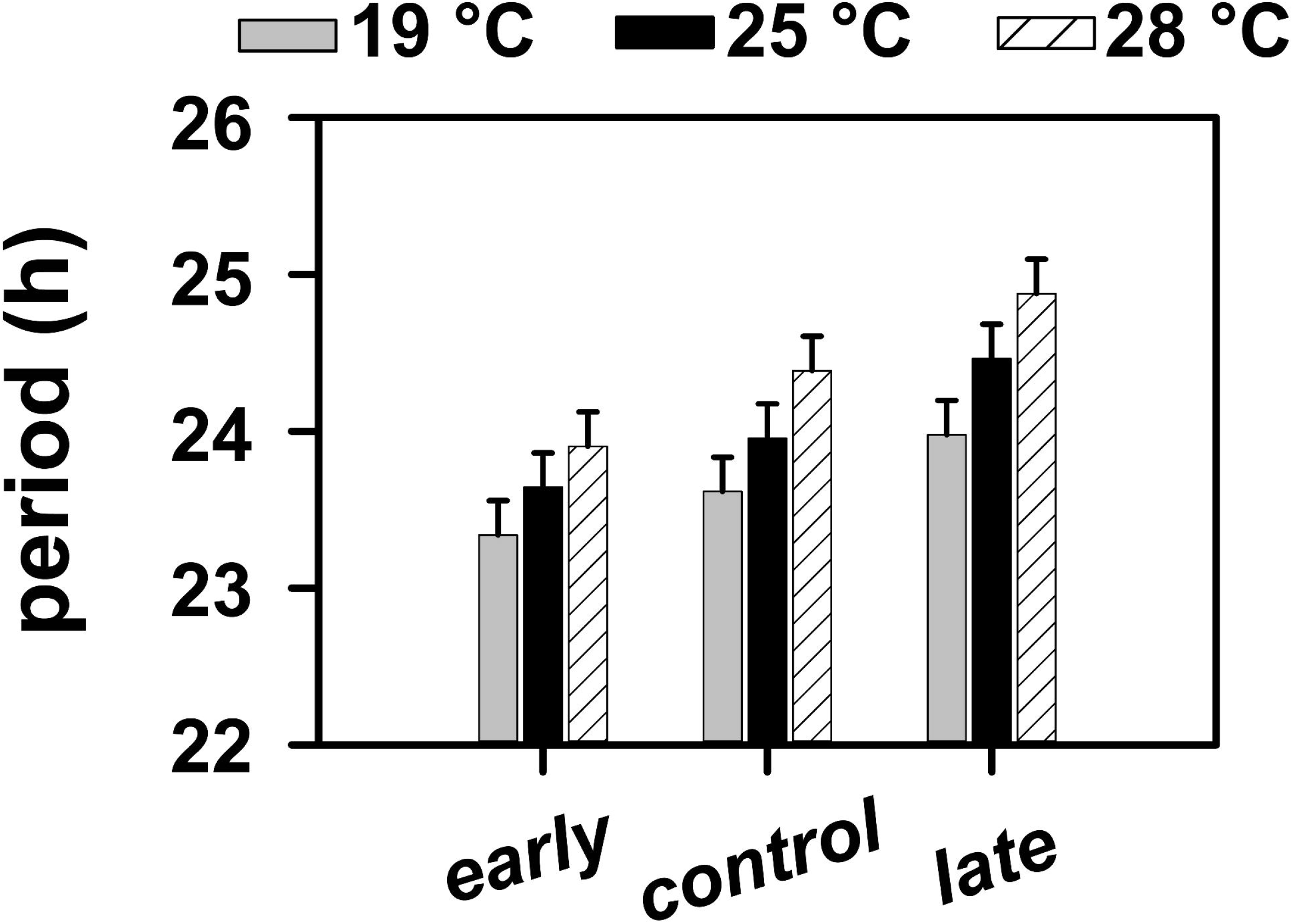
Free-running period of *early*, *control* and *late* stocks post entrainment to LD 12:12 at 19, 25 and 28 °C. All error bars are 95% CI following a Tukey’s HSD test at *α* = 0.05. Therefore means with non-overlapping error bars are significantly different from each other.

## Discussion

In this study we were interested in examining features of entrained activity/rest behaviour of *early* and *late* stocks under a variety of temperature cues in order to a) test responsiveness of the circadian clocks regulating activity/rest rhythms of these stocks to temperature cues, b) understand the mechanisms of entrainment that can account for such entrained behaviours, and c) as a consequence, understand the similarities, if any, in the organisational principles of the circuit regulating emergence rhythms and activity/rest rhythms.

An individual’s ability to re-synchronise to a new environmental schedule has been understood within the framework of the individual’s circadian clock’s phase response curve (Moore-Ede et al., 1982). A phase response curve (PRC) is a plot of phase-shifts that the individual’s clock is capable of, when stimulus of a certain strength is provided at different times of the internal cycle (Moore-Ede et al., 1982; Saunders, 2002). Therefore, for instance, if an individual’s clock’s PRC exhibits larger delay shifts than advance shifts, then said individual will re-synchronise to a westward flight across multiple time-zones much faster than an eastward flight. In the case of our *early* and *late* flies, since all stocks re-synchronise quicker to 6-h phase advances (Fig. 1) compared to delays, we infer that the temperature pulse PRC of our stocks have overall larger advance zones than delay zones. However, our results demonstrated that *late* stocks re-synchronised significantly faster to 6-h phase delay than *early* and *control* stocks (Fig. 1), thereby implying that the *late* stocks have larger delay zone than the other stocks. This would suggest the co-evolution of high amplitude temperature pulse PRCs of the circadian clock governing activity/rest rhythms in the *late* stocks in response to selection for evening adult emergence. High amplitude PRCs also imply increased phase variation (Pittendrigh and Daan, 1976), higher amplitude and power of rhythm (Brown et al., 2008; Nikhil, Vaze, et al., 2016; Vitaterna et al., 2006) and increased accuracy of entrainment (Beersma et al., 1999).

In relation to the aforementioned predictions, we obtained curious results when we analysed phases of entrainment in our stocks under TC cycles with three different durations of the thermophase. We found that while there was no between-stock difference in phases of entrainment under short thermoperiod (Fig. 2C-top-left), the *late* stocks showed significantly delayed phase compared to the *early* stocks under both, TC 12:12 and TC 18:06 (Fig. 2C-top-middle and right). In both these cases, because the *early* and *late* stocks did not individually differ from the *control* stocks, it is not possible to comment upon the individual stock’s contribution to phase lability under different temperature regimes. However, it is possible to still conclude that the small between-stock differences in lability, which may be present although not statistically detectable, could be due to between-stock differences in the PRCs.

Phase-difference between the stocks, however, can be explained using the non-parametric model of entrainment (Daan and Aschoff, 2001; Pittendrigh and Daan, 1976). The model posits that the difference between FRP and the period of the external environment is adjusted, during entrainment, via phase-shifts due to the time-cue. This response is characterised using a PRC (discussed above). Therefore, individuals with longer FRP are expected to show delayed phase under entrainment and vice versa, and this has found ample experimental evidence (Daan and Aschoff, 2001; Pittendrigh and Daan, 1976; Rémi et al., 2010; Roenneberg et al., 2005; Sharma et al., 1998; Srivastava et al., 2019; Wright et al., 2005). This however is thought to occur under the assumption that the PRC is a fixed entity in all these individuals. Therefore, if individuals have divergent PRCs they will not necessarily show such a relationship between FRP and entrained phase. We found that phases under entrainment to TC 06:18 were not different between stocks, and neither were the FRP of these stocks under constant conditions post entrainment to TC 06:18 (Fig. 2C). In case of entrainment to TC 12:12, phases of the *late* stocks were delayed and their FRP was also longer under constant darkness post entrainment (Fig. 2C). These two results are in agreement with the general rule outlined above. However, the absence of such a relationship between FRP and entrained phase under TC 18:06 reveal that although entrainment is in agreement with the non-parametric model, there is compelling support in favour of the co-evolution of divergent temperature pulse PRCs in our stocks.

To further garner support for divergent PRCs in our *early* and *late* stocks, we examined other features of entrainment to TC cycles. We found evidence for increased amplitude expansion, higher power of periodogram and higher accuracy in the *late* stocks, all of which are indicative of high amplitude PRCs. It is also interesting to note, at this point, that all the phase variations in the *late* stocks under different TC cycles is driven by the change in the evening bout of activity (see Fig. 2B). This can be clearly seen when we examined phase of evening peak of activity under all three TC cycles (Fig. 6). Importantly, analyses of the phase of evening peak of activity revealed that while the phase of *early* and *control* stocks do not differ from each other, the *late* stocks are phase delayed significantly from both, *early* and *control* stocks. This implies that plasticity in waveform in response to temperature regimes is predominantly brought about by the response of *late* stocks, which is in agreement with the idea of high amplitude temperature PRCs in these populations.

**Figure 6:**
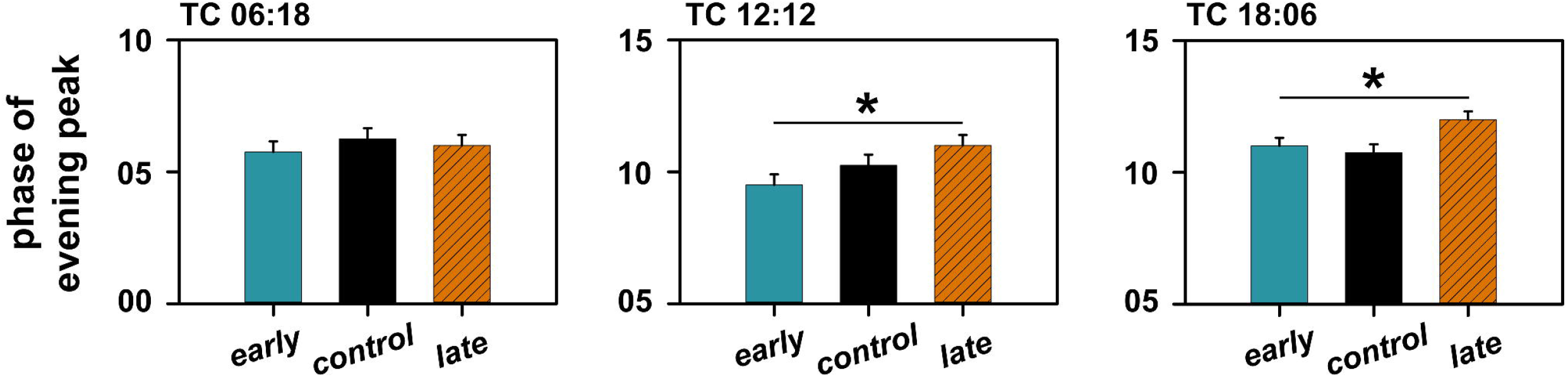
Phase of evening peak of activity of *early*, *control* and *late* under three different thermoperiods. All error bars are 95% CI from a Tukey’s HSD test at *α* = 0.05. Therefore, means with non-overlapping error bars are statistically significantly different from each other. Additionally, asterisks are drawn to indicate means that are significantly different.

Although experiments so far have indicated enhanced temperature sensitivity in the *late* stocks, we do not find strong evidence for the same under different constant ambient temperatures (Fig. 4). Previous experiments from our laboratory have shown that there is no significant difference in the photic PRCs between *early* and *late* stocks (Nikhil, Vaze, et al., 2016). Therefore, one could argue that similar extent of phase variation in the stocks across different constant temperature regimes may be a consequence of similar slopes of their photic PRCs. It is known, however, that PRC amplitudes are a function of constant ambient temperatures, such that low temperatures induce large amplitude PRCs and vice versa (Lakin-Thomas et al., 1991; Varma et al., 2013). The small differences in our stocks’ response to constant temperatures could therefore indicate small differences in the way the photic PRCs change in response to temperatures. This promises to give us a better understanding of the organisation of circadian clocks in these stocks, and may be an interesting question to pursue further. However, such small differences in the response of activity/rest behaviour in our stocks under different temperatures is in contrast with another study of ours wherein the *late* stocks show high degree of phase plasticity, albeit for the adult emergence rhythm (Abhilash et al., 2019). This we interpret as there being subtle differences in the overall organisation of the circadian network regulating emergence and activity/rest rhythms.

Plasticity of FRP under different temperature regimes pose an extremely interesting conundrum. In this regard two key results are important to note; a) FRP post entrainment to different TC cycles are shorter by > 1-h relative to the FRP post entrainment to LD cycles under different constant temperatures (Figs. 2C-bottom and 5; Table 1), and b) FRP post entrainment to short and long thermoperiod behave similarly but different from FRP post entrainment to TC 12:12 (Fig. 2C-bottom; Table 1). Temperature compensation is a crucial pre-requisite for calling an oscillatory physiological process a circadian clock (Dunlap et al., 2004; Moore-Ede et al., 1982; Saunders, 2002). This means that FRP under different constant temperatures, within physiological limits, is constant. Evidence for this comes from studies wherein under warmer temperatures, while biochemical processes should be faster, the FRP lengthens thereby indicating compensatory mechanisms (see Saunders, 2002 for examples; also Fig. 5). Such compensatory mechanisms implied that in order for entrainment to occur in response to temperature time-cues, only phase-shifts must occur (as predicted by the non-parametric model) and not period changes (as predicted by the parametric model, which posits that the zeitgeber’s effect is integrated over the cycle to constantly modulate the angular velocity of the clock, and therefore allow entrainment). However, we find that despite this being the case FRP post entrainment to different thermoperiods varies (Fig. 2C-bottom). While the period value averaged over all stocks post entrainment to TC 12:12 is ~22.98-h, FRP post entrainment to TC 06:18 and TC 18:06 lengthens and is ~23.32-h and ~23.43-h, respectively (Table 1). If these responses were due to compensatory mechanisms, one would expect opposite effects on FRP post entrainment to short and long thermoperiods. Therefore, we think that these reflect some form of after-effects due to entrainment to TC cycles. Additional support for this also comes from the result that FRP values shorten greatly after being under the influence of different TC cycles relative to values after being under the influence of LD cycles and different constant ambient temperatures. Importantly, we find that *late* stocks show significantly longer FRP post entrainment to TC 12:12, while there is no between-stock difference under the two other thermoperiods (Fig. 2C-bottom; Table 1). This suggests that the FRP of *late* stocks are less likely to change in response to temperature cycles relative to *early* stocks. While there have been many reports of after-effects of light regime on FRP (Dunlap et al., 2004), there are very few on after-effects of temperature cycles (Balzer and Hardeland, 1988). Our results, to the best of our knowledge, provides first hints of temperature after-effects on FRP in *Drosophila* activity/rest rhythms, implying that temperature may also contribute to entrainment via parametric means, warranting further, more detailed documentation of effects of temperature cycles on FRP.

In conclusion, we find enhanced temperature sensitivity of the activity/rest rhythm in *late* chronotypes. Interestingly, altered rhythm phase under different temperature regimes of the *late* stocks is driven by changes in the evening bout of activity. Further, analyses of properties of entrainment and FRP implied that results can be explained under the assumption of the evolution of divergent temperature pulse PRCs, a matter worthy of further study.

## Acknowledgements

We are very grateful to late Prof. Vijay Kumar Sharma (VKS) for providing us with the tools that enabled us to carry out this research, and for a wonderful work environment that enabled us to undertake this study. We thank Srishti Priya and Arijit Ghosh for help with experiments. We would also like to acknowledge financial support from Science and Engineering Research Board (SERB), New Delhi to VKS (EMR/2014/001188), intramural funding from Jawaharlal Nehru Centre for Advanced Scientific Research (JNCASR) and consumable grant from Department of Biotechnology (DBT), Government of India to VS (BT/INF/22/SP27679/2018).

